# Live Analysis of Position Effect Variegation (PEV) in *Drosophila* Reveals Different Modes of Action for HP1a and Su(var)3-9

**DOI:** 10.1101/2021.10.10.463835

**Authors:** Farah J. Bughio, Keith A. Maggert

## Abstract

Position Effect Variegation (PEV) results from the juxtaposition of euchromatic and heterochromatic components of eukaryotic genomes, silencing genes near the new euchromatin/heterochromatin junctions. Silencing is itself heritable through S phase, giving rise to distinctive random patterns of cell clones expressing the genes intermixed with clones in which the genes are silenced. Much of what we know about epigenetic inheritance in the soma stems from work on PEV aimed at identifying the components of the silencing machinery and its mechanism of inheritance. The roles of two central gene activities – the *Su*(*var*)*3-9*-encoded histone H3-Lysine-9 methyltransferase and the *Su*(*var*)*205*-encoded methyl-H3-Lysine-9 binding protein HP1a – have been inferred from terminal phenotypes, leaving considerable gaps in understanding how PEV behaves through development. Here, we investigate the PEV phenotypes of *Su*(*var*)*3-9* and *Su*(*var*)*205* mutations in live developing tissues. We discovered that mutation in *Su*(*var*)*205* compromises the initial establishment of PEV in early embryogenesis. Later gains of heterochromatin-induced gene silencing are possible, but are unstable and lost rapidly. In contrast, a strain with mutation in *Su*(*var*)*3-9* exhibits robust silencing early in development, but fails to maintain it through subsequent cell divisions. Our analyses show that while the terminal phenotypes of these mutations may appear identical, they have arrived at them through different developmental trajectories. We discuss how our findings expand and clarify existing models for epigenetic inheritance of heterochromatin-induced gene silencing.

**Significance:** Current concepts of epigenetic inheritance are exemplified by Position Effect Variegation, a phenomenon whereby heterochromatin can repress genes in clonal cell lineages. Heterochromatin is required for genome protection as it silences toxic transposable elements and prevents instability of repeat sequences. Histone H3 modified by methylation of Lysine-9 and HP1a are critical components of heterochromatin. Using live cell analysis of PEV in mutants in strains with mutations in *Su*(*var*)*3-9*, which encodes the histone methyltransferase, and *Su*(*var*)*205*, which encodes HP1a, we describe an unexpected dynamism in PEV, challenging current models of epigenetics, and revealing unexpectedly different modes of action of these two fundamental components of heterochromatin.

## Introduction

Position Effect Variegation (PEV) was first observed in *Drosophila* as random “eversporting” patterns of *white^+^* gene expression in individual ommatidia of the compound eye (1). Genes undergoing PEV did so because genome rearrangements (*e.g*., chromosome inversions, transpositions) created new heterochromatin-euchromatin breakpoints (2). Rearrangements brought normally-euchromatic genes into juxtaposition with heterochromatin and normally heterochromatic-genes with euchromatin (3–5). In both cases, genes near those rearrangement breakpoints were repressed in some cells but in others they were not, resulting in random and compelling patterns of expression in otherwise genetically-identical cells.

The current view of PEV derives mostly from work on heterochromatin-induced gene silencing of euchromatic reporter genes placed near heterochromatin (6). From these, it is envisioned that heterochromatin proximity induces silencing on the reporter gene when heterochromatin forces the acquisition of methylation of Lysine-9 of Histone H3 (H3K9) near the breakpoint including in those nucleosomes that package the reporter gene. A necessary component of this view is that heterochromatin-induced gene silencing “spreads” from the heterochromatin to juxtaposed DNA, regardless of its sequence, bringing heterochromatin’s intrinsic repression to closely-linked genes. Close analysis of H3K9 methylation status on genes linked to breakpoints confirm an increase in H3K9 methylation on them, however the “spreading” appears discontinuous, focused at gene promoters (7–8). The manifest silencing itself is also discontinuous, occasionally “skipping over” one or more genes (9–10). These studies indicate that the phenomenon of PEV is best envisioned as multiple separable phenotypes – the mechanism of silencing brought about by heterochromatin, the degree to which “spreading” may occur, the extent of spreading from the heterochromatin source, and whether/how a gene can escape silencing (11).

Because the patterns of gene silencing in each cell are inherited through cell division, leading to the classical clonal patterns of expression in PEV, it seems clear that PEV requires an epigenetic memory of the extent of heterochromatin spreading. In fact, spreading itself demands it to be so. Epigenetic memory must either encode the distance of spreading every cell generation, or it must encode the particular modifications on each heterochromatic nucleosome after spreading first occurs.

Genetic screens for genic *Suppressors of* [Position Effect] *Variegation* (*i.e*., *Su*(*var*)s, second-site mutations that alleviate the silencing of heterochromatin, returning a variegating allele to a more wild-type expression state), uncovered many loci that have been found to encode major protein components of heterochromatin (12). The gene product of the *Drosophila Su*(*var*)*3-9* locus is one of three Histone Methyltransferases – along with *G9a* and *eggless/ SETDB1* – capable of methylating Histone H3K9. The H3K9 methylation recruits Heterochromatin Protein 1 (HP1a), originally identified cytologically and later found to be encoded by the *Su*(*var*)*205* locus (6). It is HP1a that effects gene silencing. Mutations in either *Su*(*var*) gene derepress genes being silenced by heterochromatin, and in fact these are the two strongest genic modifiers of PEV yet-described, restoring expression of most alleles to wildtype.

The accepted model for PEV and the roles of *Su*(*var*)*3-9* and *Su*(*var*)*205* are based on dominant loss of function phenotypes in terminally-differentiated tissues, almost always the expression of the *white^+^* gene in adult ommatidial pigment cells or the expression of the *yellow*^+^ gene in adult bristles or abdominal cuticle. However, there are many trajectories that silencing can take through the manifold cell divisions and differentiations during development before arriving at an endpoint in adult organisms. We were inspired by Dr. Janice Spofford’s review of PEV from 1976 (2), *“In flies, the mosaic phenotypes* [PEV] *are not expressed until the final cell divisions; descendants of known single cells have not been sampled sequentially during the history of a cell lineage, and the degree of reversibility of the inactive state and the consequent identification of the time of inactivation remain moot.”* Live monitoring of PEV throughout development would help resolve the times at which silencing is set, when it is lost, and when *Su*(*var*)*205* and *Su*(*var*)*3-9* act on PEV.

We recently developed the Switch Monitoring (SwiM) System (Figure 1A) to monitor gains and losses of heterochromatin-induced gene silencing in a live PEV model in *Drosophila* (11). For the SwiM System, we embedded a ubiquitously-expressed gene encoding GAL80, the yeast GAL4-specific transcriptional repressor, in heterochromatin causing it to undergo PEV. Simultaneous use of a ubiquitously-expressed yeast GAL4 transactivator allowed us to monitor GAL80 PEV by assaying GAL4 activity (*i.e*., GAL4p in the absence of GAL80p). We employed the dual fluorescent reporter *G-TRACE* lineage tracing system (13) to identify those cells in which GAL80 was repressed, expressed, or those in which GAL80 had undergone rounds of gains or losses (“switches”) of heterochromatin-induced gene silencing. The SwiM System proved to be remarkably rich, as we could infer the histories of switching in individual cell clones/lineages by analyzing the presence/absence and intensities of GFP and RFP fluorescence from the *G-TRACE* component (Figure 1B), the size of like-expressing clones, the fluorescence patterns within clones, and the proximity of clones to those of other expression patterns (11). Using SwiM we previously concluded that PEV is highly dynamic through development, specifically that Spofford’s *“reversibility of the inactive state”* was very high, recapitulating analyses by Eissenberg and colleagues (14–15). Through SwiM system analysis we also detected a *reversibility of the active state*, for the first time showing that heterochromatin-induced gene silencing could be reacquired once lost [although see (16)], arguing that the stable maintenance of heterochromatin-induced gene silencing is not simply established early in development and lost by defect in the epigenetic memory mechanism. Rather, as Spofford predicted, we could infer rich and varied trajectories of silencing toward the final “mosaic” phenotypes of PEV not expressed until the final cell divisions.

**Figure 1.**
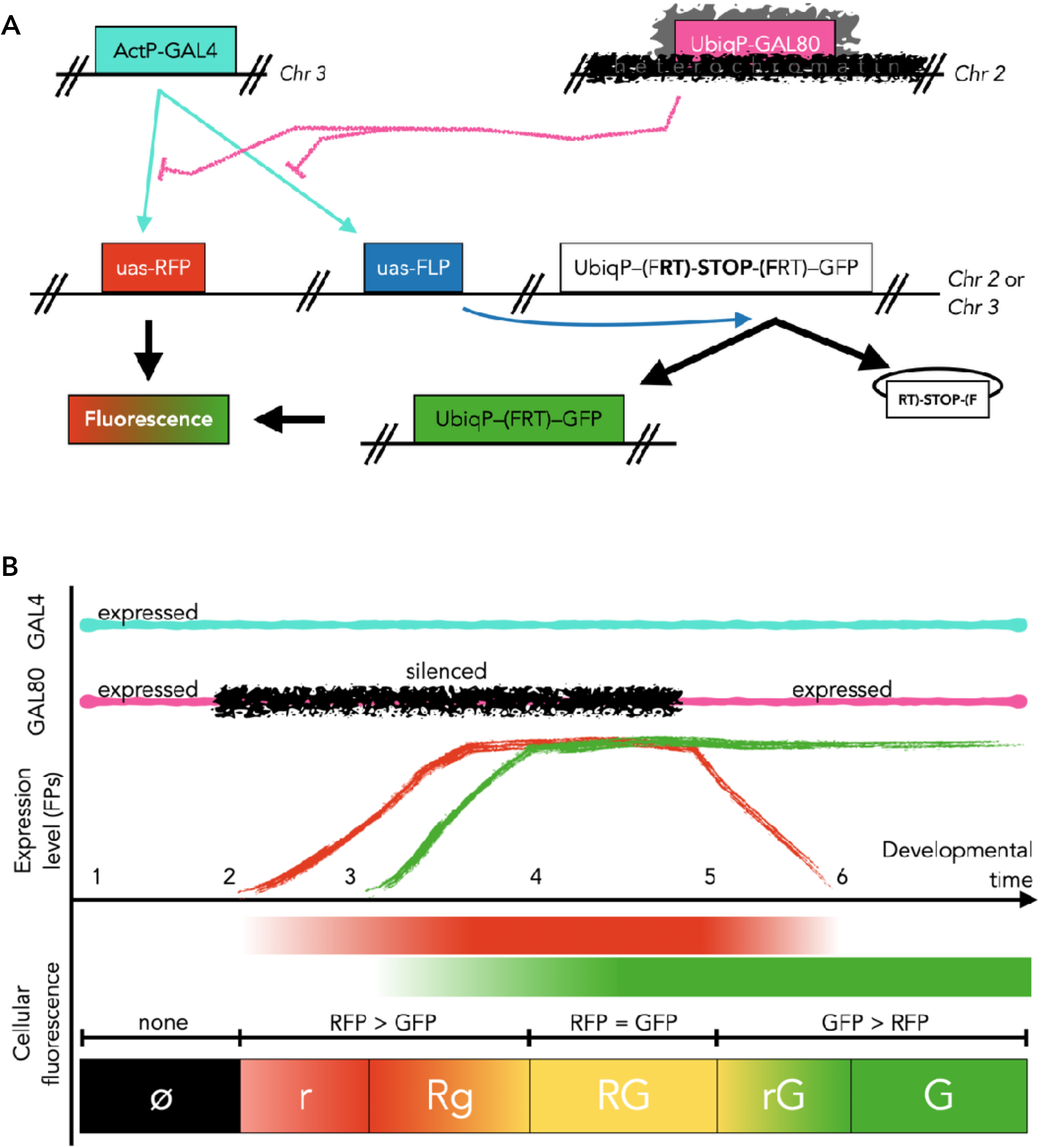
Schematic of the Switch Monitoring (SwiM) System. **(A)** The SwiM System includes ubiquitously-expressed GAL4 activator (teal “ActP-GAL4”) and GAL80 repressor (pink “UbiqP-GAL80”) genes, the latter transposed into the heterochromatin under study. GAL4 activity in the absence of GAL80 activity is monitored by the multi-component *G-TRACE* lineage tracer (13). GAL4 transactivates a UAS-RFP (red “uas-RFP”) transgene directly, and thus reports on repression of the GAL80 gene. GAL4 also transactivates the FLP site-specific recombinase (blue “uas-FLP”) which catalyzes the removal of a STOP cassette and permanent ubiquitous activation of GFP [converting the inactive white “UbiqP-(FRT)-STOP-(FRT)-GFP” to the active green “UbiqP-(FRT)-GFP” and the “RT)-STOP-(F” extrachromosomal circle, which is lost through mitosis]. **(B)** Expression of GAL4 (teal) and switches in silencing of GAL80 (pink and black) lead to RFP and GFP fluorescence. The comparison of RFP and GFP fluorescence levels are indicative of whether silencing is intact, compromised, or has switched. 1-6 indicate hypothetical developmental timepoints that are discriminated by the RFP/GFP fluorescence levels. “ø” indicates no fluorescence; “r” indicates low RFP fluorescence with no GFP fluorescence at the onset of GAL80 silencing; “Rg” occurs later as RFP reaches maximal levels and GFP begins to build up; “RG” indicates robust expression during times in which GAL80 is silenced; “rG” indicates that silencing has been lost and the RFP is decaying while GFP expression persists; “G” indicates that silencing is no longer existent leaving just the persistent GFP.

Since the accepted model of PEV is based largely on the phenotypes and biochemical activities of Su(var)3-9 and HP1a, namely the continuing necessity of their presence to maintain silencing in somatic cells, it became important to analyze mutants using the SwiM System. In this work, we do so. We extend our previous work on PEV by showing that the strongly suppressive *Su*(*var*)*3-9*^1^ mutation of the histone methyltransferase and the *Su*(*var*)*205*^5^ mutation of the methyl-histone binding protein exert their effects on PEV at distinct times in development. These alleles are commonly used to investigate the roles of these proteins in studies of PEV. In this work, we show that SwiM System analysis indicates heterozygous *Su*(*var*)*205*^5^ mutants exhibit an almost complete loss of silencing in early embryogenesis, and the lack of silencing seems to persist until adulthood. We observed that individual cells rarely acquired silencing, and those that did lost it again probably within the same cell cycle. We could only detect these rare events indirectly, in contrast to *wild-type* organisms where gains of silencing are relatively common. In contrast to *Su*(*var*)*205*^5^, heterozygous *Su*(*var*)*3-9*^1^ mutants were able to well-establish silencing early in development, but we observed a progressive loss of silencing through development. Specifically, even at late stages, mutants of *Su*(*var*)*3-9* were better able to establish silencing, but it was nonetheless rapidly lost. Analysis of both mutant conditions is consistent with our developing idea that the balance between relative stability of expression state and occasional switches leading to PEV may be mediated by changes in nonchromosome-bound factors – for example HP1a protein level – that differ between cells and cell clones.

## Results and Discussion

### The expressivity of PEV is a wide spectrum

The GAL80 component of the SwiM System contains a *white^+^* transgene that can be scored in adult eyes, which allowed us to easily estimate the variation of silencing occurring at the heterochromatic GAL80/*white*^+^ locus within a population of isogenic organisms. We measured the amount of pigmentation in 104 *w/Y*; *P*{*white*^+mC^=*tubP-GAL80*^ts^}^10-PEV-80.4^/+ adult eyes, dependently scoring both left and right eyes, and dividing them into categories of variegation by quantifying the number of ommatidia expressing *white^+^* (Figure 2). We observed that about one-third of the flies had near-absent *white^+^* pigmentation (“0-20” category), indicating robust heterochromatin-dependent gene silencing (Figure 2A, D). The remainder had intermediate levels of silencing and expressed classical “Patched” patterns of *white^+^* PEV (Figure 2B), and few flies (~5%) had a high level of *white^+^* pigmentation (Figure 2C), indicating organisms with weak heterochromatin-induced gene silencing at the GAL80/*white*^+^. Although we did not directly test GAL80 and *white^+^* variegation in the same individuals, our previous work with *white^+^* and a closely-linked *GFP* showed that variegation of closely-linked (*i.e*., within the same *P-element* transposon) is well-correlated (11). Wang and Elgin have observed a remarkable reproducibility in PEV expressivity of specific variegating alleles after multiple generations of selection (17). In the variegation of the SwiM System GAL80 allele, we could detect no such reproducibility, nor could we detect any influence by unlinked modifiers of variegation in our strains (11). The difference between their observations and the behaviors of the GAL80 locus we use here is not immediately clear, however it may be that they selected for expression in regions of the eye that poorly repress variegating gene expression (18), or for modifiers that have preferential effects at certain loci (19).

**Figure 2.**
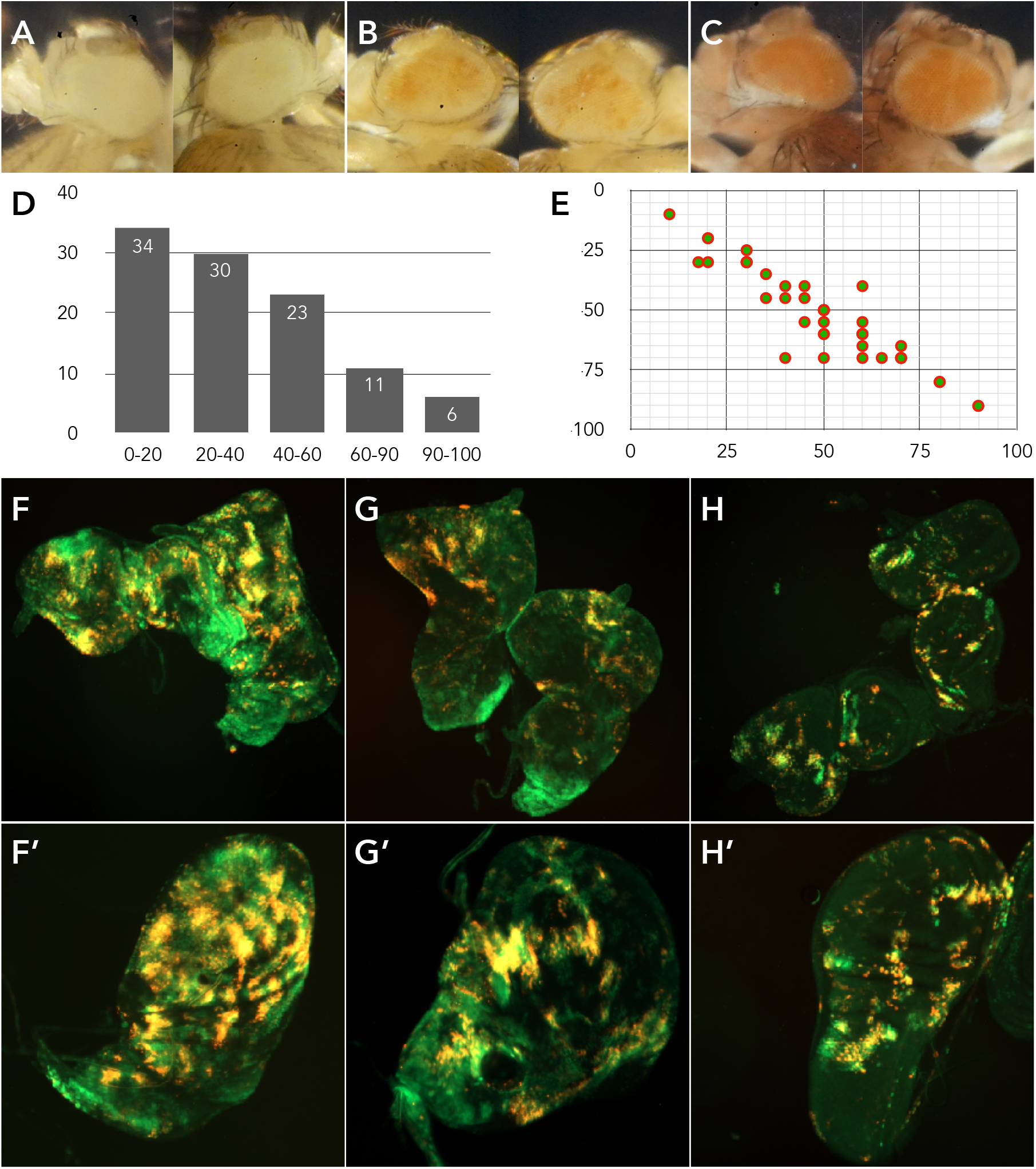
Extent of PEV (silencing) is correlated within individuals, but variant within populations. **(A-C)** Paired left and right eyes of representative categories of PEV expressed by *w/Y;* +/*P*{*white*^+mC^=*tubP-GAL80*^ts^}10-PEV-80.4 flies, showing that silencing within an individual affects the entire organism similarly (11). (A) shows complete silencing (*i.e*., “0” expression), (B) shows mid-level (“40-60%”), and (C) shows near *wild-type* expression (“90-100%”). **(D)** Histogram of data taken from scoring 52 *w*/*Y*; +/*P*{*white*^+mC^=*tubP-GAL80*^ts^}10-PEV-80.4 flies. **(E)** Dissected eye and wing imaginal discs were categorized for expression, scoring the fraction of cells expressing GFP, RFP, or the overlap within each imaginal disc. Average fluorescence categories of eye discs (X-axis) correlate well with average fluorescence categories of wing discs (Y-axis) from the same individual. Non-parametric analysis cannot discriminate any difference between eye and wing fluorescences, indicating they come from indiscriminable populations (χ^2^ = 51.1; *P* = 0.43). Non-parametric regression was close to unity (Kendall’s robust line-fit, *b* = 0.91). **(F-H)** Images of representative eye-antennal imaginal discs (F-H) and wing discs (F’-H’) taken from the same organisms. (F) shows robust silencing of the GAL80 component of the SwiM System, (G) shows medial level, and (H) shows poor silencing.

The high variability in studies of PEV makes many assessments fraught: proportions of cells experiencing silencing versus expression vary too much between individuals to produce robust or meaningful statistical descriptions, and population measurements (*e.g*., pigment extractions) lose meaningful information (*e.g*., the patterns of expression). One of the pronounced benefits of the SwiM System is the ability to infer individual trajectories of silencing through development by analyzing whole organs (*i.e*., imaginal discs) from individuals, and thereby more-completely understand the breadth of histories of heterochromatin-induced gene silencing in cell clones as they expand.

In contrast to the pronounced variability between individuals, the variability in silencing within individuals was minimal (Figure 2E; examples of high, medium and low silencing are shown in Figures 2F-F’, G-G’, and 2H-H’, respectively). We individually dissected eye/antennal and wing imaginal discs and compared the degree of silencing and expression as revealed by the SwiM System. We found a strong correlation between degree of expression or silencing in all four discs of individuals, similar to our previous results showing good organism-wide correlation of heterochromatin-induced gene silencing (11). This validates the use of SwiM to understand silencing through development, and extends our analysis of eye discs in our previous work to wing discs in this work. By shifting to SwiM System analysis of wing discs we abrogate concerns about the influence of the eye-specific enhancer/promoter combination of the *mini-white* (+*mC*) gene in the *P*{*white*^+mC^*=tubP-GAL80*^ts^}^10-PEV-80.4^ component of the SwiM System.

### Switch Monitoring System analysis of Position Effect Variegation in *wild-type* organisms

We analyzed the *wild-type* patterns of heterochromatin-induced gene silencing in multiple wing imaginal discs dissected from +/*P*{*white*^+mC^*=tubP-GAL80*^ts^}^10-PEV-80.4^; *G-TRACE*/ *P*{*white*^+mC^=*Act5C-GAL4*}*17bFO1* animals. The patterns provided us a baseline to understand how establishment and maintenance of silencing manifest in comparison to mutations in *Su*(*var*)*205* and *Su*(*var*)*3-9* animals, which are expected to appear as differences from these patterns.

In organisms *wild-type* for components of heterochromatin, expression of RFP and GFP in fluorescent clones fell broadly into three categories (Figure 3A-B). First, clones expressing bright RFP and GFP fluorescence (the overlap appearing as bright yellow fluorescence) indicated cells in which silencing of GAL80 was ongoing (Figure 3C-D). In these cells, the absence of GAL80 due to silencing allowed the GAL4-dependent RFP to express, and the GAL4-dependent *FLP*-dependent rearrangement to permanently activate the GFP transgene. The relative brightness of RFP and GFP indicated that the silencing of GAL80 had been ongoing long enough to allow both RFP and GFP accumulation, corresponding to Figure 1B, between developmental time points 4 and 5, “RG” type. It is possible that some of these cells had very recently lost heterochromatin-induced gene silencing at the GAL80 locus because the perdurance of RFP is on the order of 6-8 hours, which defines the temporal resolution of our Switch Monitoring analyses (11).

**Figure 3.**
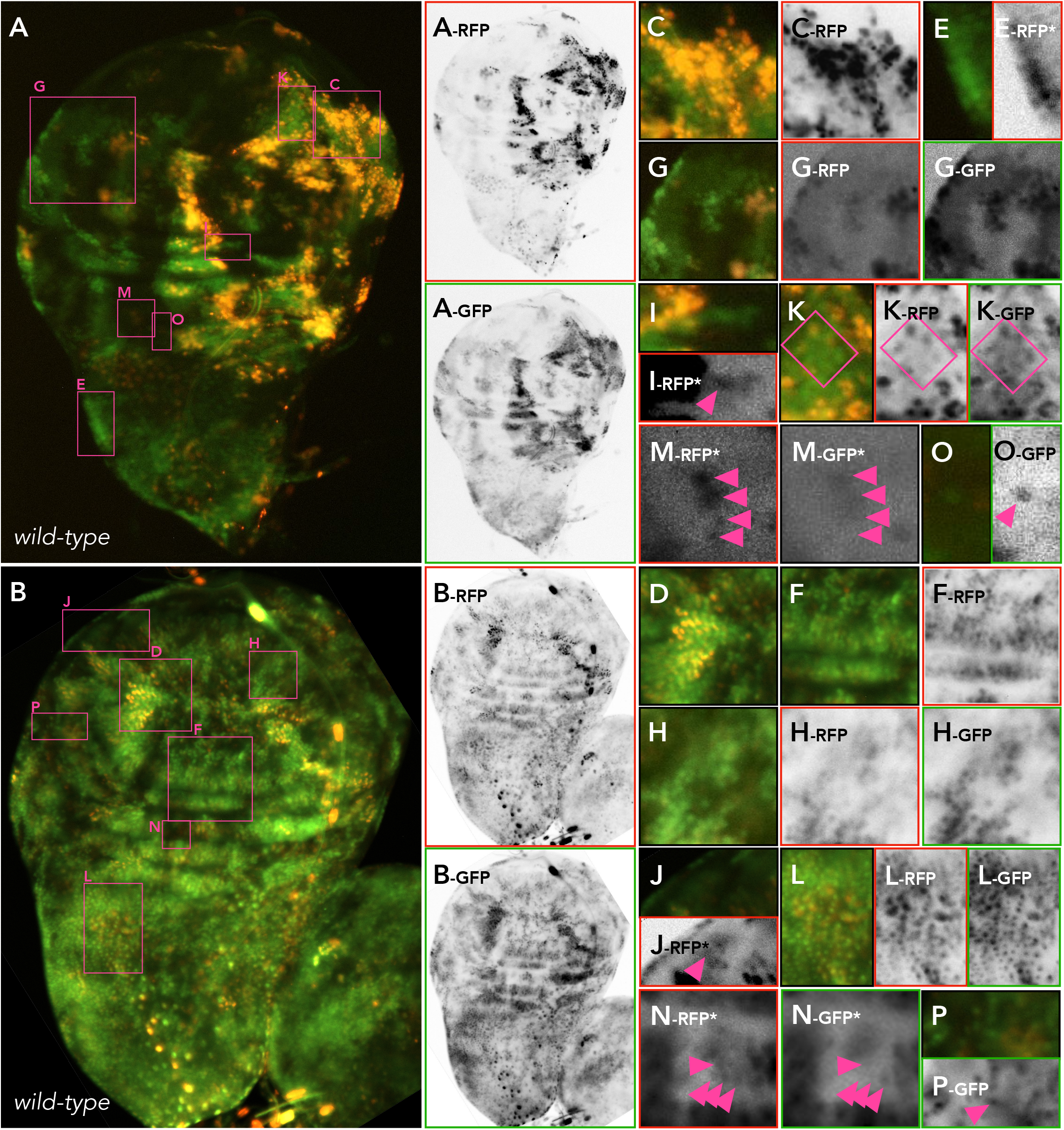
Switch Monitoring System analysis of PEV in wild-type organisms. **(A-B)** Wholemount wing imaginal discs showing RFP and GFP fluorescence as revealed by SwiM System analysis of +/*P*{*white*^+mC^=*tubP-GAL80*^ts^}^10-PEV-80.4^; *G-TRACE*/*P*{*white*^+mC^=*Act5C-GAL4*}*17bF01* flies. A-rfp, A-gfp, B-rfp, and B-gfp show inverted monochrome separations of the red and green color channels. Images in (A) and (B) are two discs taken from separate individuals and are both presented and analyzed to show the common features of the patterns revealed by SwiM System analysis. Pink boxes indicate location of enlarged regions shown in (C-P). **(C-D)** show regions of imaginal discs in which silencing of GAL80 is ongoing, appearing as robust RFP and GFP fluorescence. **(E-F)** show regions that have undergone relatively recent losses of silencing of GAL80, evident by diminished RFP and robust GFP. (E-rfp*) shows the inverted monochrome RFP separation of (E), the asterisk (*) indicating that the image has been adjusted for bright-contrast to reveal low levels of RFP. **(G-H)** show regions that, like (E-F), have lost silencing and express GFP but not RFP. These regions must have lost silencing relatively long before observation as RFP has had time to decay. **(I-J)** show regions that, like (G) and (H), have no obvious RFP. However the inverted monochrome RFP separations, (I-rfp*) and (J-rfp*), reveal very low levels of RFP in some cells (triangles). **(K-L)** show regions regions intermixed with bright and dim RFP within fields of uniform GFP. **(M-N)** show individual cells expressing RFP and no detectable GFP. These represent cells that have recently acquired gene silencing of the GAL80 gene (triangles). **(O-P)** show isolated GFP-expressing cells, as confirmed by the (O-gfp) and (P-gfp) inverted monochrome GFP separations.

Second, there were many clones of cells which express GFP and some low level of RFP (Figure 3E-F). We interpret these to be cells which had sufficient heterochromatin-induced gene silencing of GAL80, followed afterward by a loss of that silencing. This first allowed GAL4-dependent activation of RFP and GFP, but the following silencing of GAL80 caused the loss of RFP expression (Figure 1B, between developmental time points 5 and 6, “rG” type). In these cells, the loss of silencing must have happened recently relative to observation, since the diminishing RFP had not degraded entirely by the time of observation.

Third, we observed some contiguous clones in which individual cells expressed GFP and no detectable RFP (Figure 3G-H). These were cells in which silencing had been established, as in the second category, but was then lost. Enough time had then passed to allow full derepression of GAL80, GAL80’s subsequent repression of GAL4, and decay of GAL4-dependent RFP mRNA and protein (Figure 1B, developmental time point 6, “G” type). We find support for this interpretation because over-manipulation of bright/contrast often revealed a very small amount of RFP fluorescence in some of these cells (Figure 3I-J).

Apart from these three broad categories of clones, there were cells that appeared to be clonally-related that were intermixes of the first and second fluorescent phenotypes (Figure 3K-L). The lineage-tracing properties of the SwiM System indicate a possible life-history for the cells in these clones. We interpret that a single ancestor cell, or multiple cells in the same clone, established silencing of GAL80 (causing them to express RFP and GFP), then silencing was lost in a subset of cells (switching those cells to GFP alone) as the clone continued to expand. We note that the intensity of GFP in these were more-or-less equal, further supporting our interpretation that the initial establishment of silencing was a single or synchronous event much earlier in the lineage. In contrast, the variation in RFP indicates asynchrony in the subsequent loss of silencing.

Some cells expressed RFP alone, and often these cells were near each other (Figure 3M-N, also squat triangle in Figure 3H), as we have commented on previously (11). Based on SwiM System analysis, these indicate cells in which silencing of GAL80 had not been established – in early embryogenesis or the first two larval instars – until within only the last few hours prior to observation (Figure 1B, developmental time point 2, “r” cells). We interpret these cells to be evidence of gains of heterochromatin-induced gene silencing. Related to this ongoing instability, those clones that are GFP, or occasionally lone GFP-expressing cells within non-fluorescent clones (Figure 3O-P), indicate that individual cells may gain and subsequently lose silencing while their cousin cells remain persistently un-silenced. Cells of this type were mentioned by Ahmad and Henikoff (16), although their assay system did not allow them to assertively conclude a definitive gain of silencing.

Finally, non-fluorescent clones or cells (Figure 3A-B and, for example, those cells in Figure 3M-P that are not indicated) can only come from complete and ongoing failure to silence GAL80 because, if GAL80 were to be repressed at any time in development, the cell and its descendent should be permanently labelled by GFP expression. This category of cells – present but not common in *wild-type* – is the one we expected to find more frequently in conditions compromising the establishment or maintenance of heterochromatin-induced gene silencing.

### *Su*(*var*)*205*^5^ mutation, reducing HP1a function, impacts early establishment and later maintenance

To understand the developmental impact of reduced HP1a function on PEV, we created flies heterozygous for the *Su*(*var*)*205*^5^ allele in conjunction with the Switch Monitoring System. This allele is a frame-shift mutation after 10 amino acids (20) and is likely amorphic. *Su*(*var*)*205*^5^ was introduced through the mother to reveal both maternal and zygotic gene product requirements (11, 14–15, 21). We dissected and analyzed *Su*(*var*)*205*^5^/*P{white^+mC^=tubP-GAL80*^ts^}^10-PEV-80.4^; *G-TRACE*/*P*{*white*^+mC^=*Act5C-GAL4*}*17bFO1* third instar larval imaginal discs.

For the most part, we saw a dramatic expansion of non-fluorescent clones relative to *wild-type*, covering the majority of every disc that we analyzed (Figures 4A-B). These cell clones had persistent GAL80-mediated repression of the GAL4-controlled components of the *G-TRACE* lineage marker, indicating that heterochromatin-induced gene silencing of GAL80 had never been experienced in most cells of the organism. This includes the two particularly critical phases of heterochromatin function, one early acting (peri-fertilization, discussed at length by us (11)) and one later (peri-gastrulation, identified by Eissenberg (15)). Both of these phases were strongly compromised in *Su*(*var*)*205*^5^ heterozygotes, but so was silencing outside of these phases, indicating that *Su*(*var*)*205*^5^ mutation has effects throughout development and not just at specific stages of tissue determination and differentiation (20). Our findings here are therefore consistent with others’, identifying HP1a as a dose-dependent structural component of heterochromatin function in early embryogenesis.

**Figure 4.**
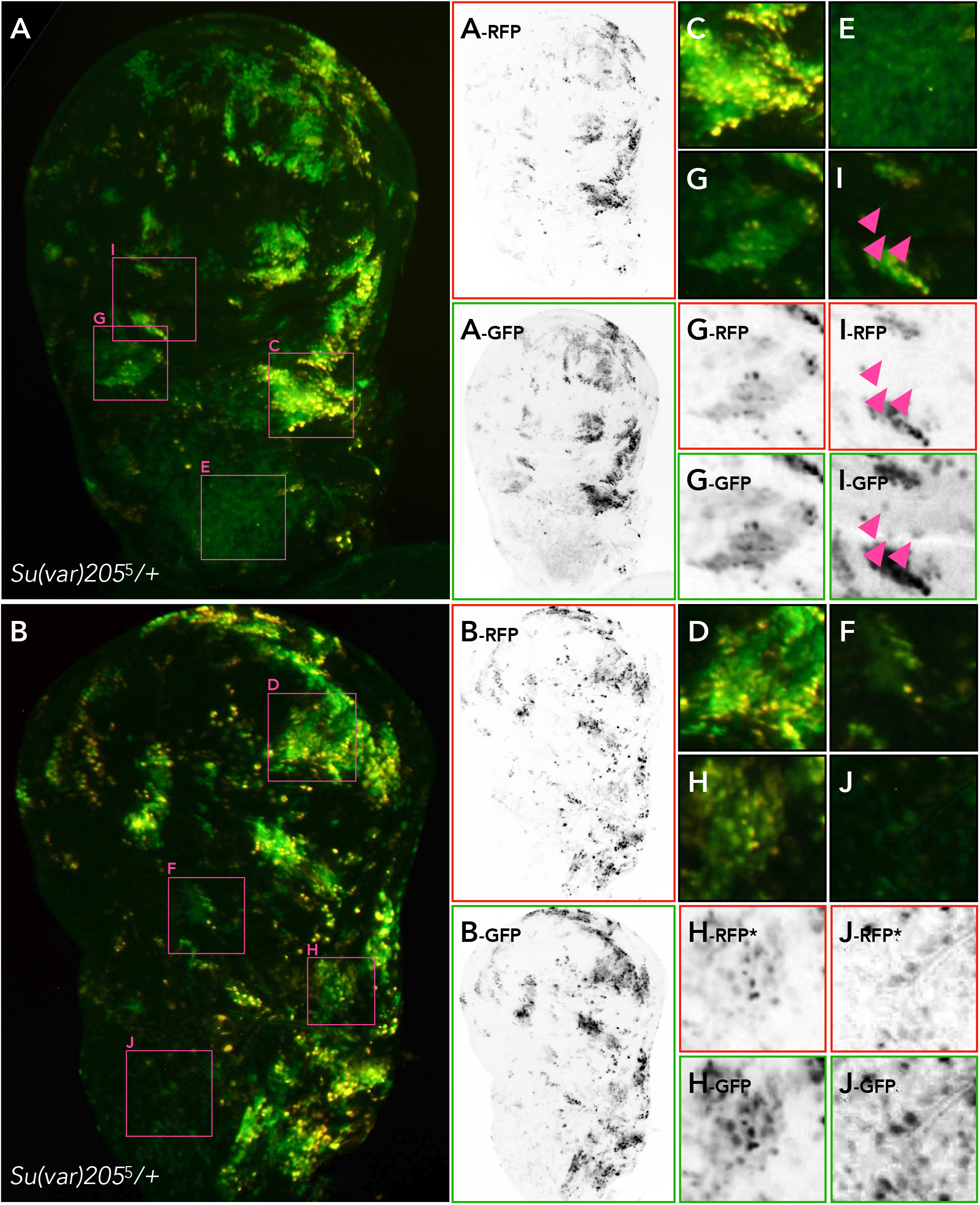
Switch Monitoring System analysis of PEV in *Su*(*var*)*205^5^/+* organisms. **(A-B)** Whole-mount wing imaginal discs showing RFP and GFP fluorescence as revealed by SwiM System analysis of genotype *Su*(*var*)*205*^5^/*P*{*white*^+mC^=*tubP-GAL80*^ts^}^10-PEV-80.4^; *G-TRACE/ P*{*white*^+mC^=*Act5C-GAL4*}17bF01. A-rfp, A-gfp, B-rfp, and B-gfp are inverted monochrome separations of the RFP and GFP color channels. Images in (A) and (B) are two discs taken from separate individuals and are presented and analyzed to show the common features of the patterns revealed by SwiM System analysis. Pink boxes indicate location of enlarged regions shown in (C-J). **(C-D)** show regions with GFP and RFP expression, indicating ongoing silencing of GAL80. Note in these regions that the variability of GFP and RFP is higher than in discs from *wild-type* organisms (Figure 3). **(E-H)** show regions with variable GFP expression and little RFP within single clones. Those in (E) and (F) were presumably earlier in their gains and losses of silencing, while those in (G) and (H) were more recent in their gains and losses of silencing. **(I-J)** show individual cells expressing GFP but not RFP (triangles). Such cells indicate ongoing variation, here thought to be short-lived gains of silencing followed by almost-immediate loss.

Despite the mutation of *Su*(*var*)*205* and the dramatic decrease in most cells’ ability to establish repression, some cells were capable of both establishing and maintaining silencing. These GFP- and RFP-expressing cells were much-less numerous in *Su*(*var*)*205*^5^ heterozygotes, but were nonetheless consistently observable in analyzed discs (Figure 4C-D). As in *wild-type* discs, the cells capable of continued silencing were generally found near each other in clones. Also as in *wild-type*, these robustly RFP-expressing cells were intermixed with GFP-expressing cells that had no or low levels of RFP.

Notably, the levels of GFP were variant within these clones. These variances could arise from asynchrony in division of cells within the clones (leading to dilution of GFP through mitosis), or may indicate that the onset of GFP expression was not synchronous as it would be if it had occurred in a single cell or at a specific time in development. The latter interpretation would indicate that cells within these clones gained silencing throughout development, during or after clonal expansion, but then later lost that silencing. We favor this interpretation because the variability we observe is higher in *Su*(*var*)*205*^5^/+ mutations than in *wild-type* (here an in (11), and from other evidence of ongoing instability.

We also interpreted clones with uniformly-brighter and uniformly-dimmer GFP fluorescence, both with low RFP fluorescence (Figure 4E-F and 4G-H), to be evidence for clones with a propensity to lose any acquired silencing. We believe the brighter clones to be cells descended from ancestors that experienced one or few early gains of silencing followed promptly by complete loss of silencing, permanently tagging all descendent cells with GFP despite the absence of any perduring RFP. We believe the dimmer clones to have had a similar history, although having happened more recently, after the clone had expanded. Without a clear coupling of silencing and developmental stage, it seems likely that the variation we see in GFP and RFP expression indicate that, as these clones develop, they undergo frequent switching, gaining and losing silencing repeatedly. Consistent with this, we could find individual cells within the latter type of clone that were both RFP-expressing and brighter GFP-expressing, although we cannot rule out that those cells have undergone a now second round of silencing. The presence of GFP-expressing cells with no detectable RFP further shows that silencing may be established in *Su*(*var*)*205*^5^ heterozygotes, but with an increased likelihood of losing that silencing. This is generally evident in comparison to discs from *wild-type* organisms (*cf*. Figure 3) which had more uniformly bright GFP.

From these behaviors, we can predict that gains of silencing should be rare or nonexistent. As we showed in previous work (11), recent gains are easily detected as RFP expression without any detectable GFP expression (Figure 1B, r phase). We could not identify any cells within otherwise non-fluorescent clones that had detectable RFP that were not also brightly-expressing for GFP. We did, however, note some isolated GFP cells within non-fluorescent clones (Figure 4I-J). We interpret these as cells that were silenced for a short enough time so that RFP could not rise to a detectable level, but nonetheless expressed sufficient FLP to catalyze the permanent activation of ubiquitous GFP expression. These lone GFP cells, as far as we can ascertain, are unique to this genotype.

Most fluorescent cell types indicate a high level of instability of heterochromatin-induced gene silencing, thus we believe that *Su*(*var*)*205*^5^ mutation acts by rendering cells less capable of establishing silencing *and* less capable of maintaining it. These defects in establishing silencing – non-fluorescent cells from ongoing defects starting in early embryogenesis, and green fluorescent cells from defects any time henceforth – are responsible for the well-characterized near-complete suppression of PEV in adult tissues. The SwiM System, however, allows us to clearly delineate very different life histories of these cells on their way to an ultimately common phenotype.

Within the cells that could, at least sporadically, establish silencing, there were clear differences in the variation of intensities for RFP and GFP when comparing *Su*(*var*)*205*^5^ mutant and *wild-type* clones. The former had higher variation despite the clone sizes being smaller. This condition is predicted if, in *Su*(*var*)*205*^5^ mutants, cells are more likely to switch from silenced to non-silenced *and* from non-silenced to silenced more frequently. We proposed in our previous work that such rapid switches might underlie the overall suppression by mutations in *Su*(*var*) genes. We also proposed that rapid switches are responsible for the types of patterns of adult PEV – clonal patches versus salt-and-pepper variegation. Under this model, large clonal patches of expression would be the result of lower instability (lower switch rates leading to larger clones of consistently-acting cells), while the smaller salt-and-pepper spots of PEV would be the result of higher instability (higher switch rates leading to smaller clones with higher variability between cells) (11).

### *Su*(*var*)*3-9*^1^ mutation, reducing a key heterochromatic histone methyltransferase, preferentially affects maintenance of PEV

Heterozygous loss-of-function mutants of the *Su*(*var*)*3-9* gene, which encodes one of the three *Drosophila* H3K9 histone methyltransferases (HMT), largely do not exhibit a loss of the initial establishment of heterochromatin-induced gene silencing. This is evident from the overall expression of GFP (Figure 5A-B) in discs taken from *G-TRACE*/*P*{*white*^+mC^=*tubP-GAL80*^ts^}^10-PEV-80.4^; *Su*(*var*)*3-9*^1^/*P*{*white*^+mC^=*Act5C-GAL4*}*17bFO1* individuals.

**Figure 5.**
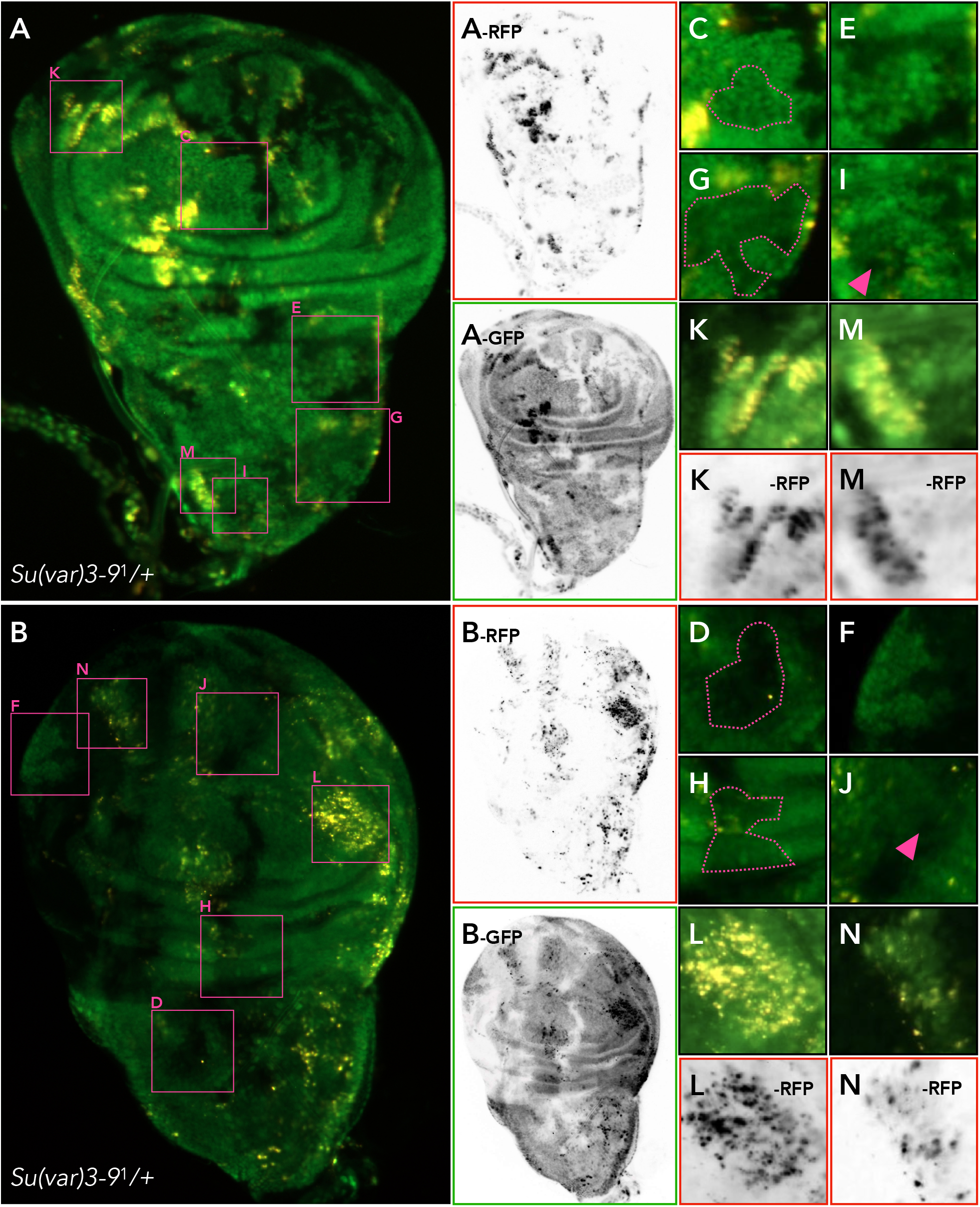
Switch Monitoring System analysis of PEV in *Su*(*var*)*3-9^1^/+* organisms. **(A-B)** Whole-mount wing imaginal discs showing RFP and GFP fluorescence as revealed by SwiM System analysis of genotype *G-TRACE*/*P*{*white*^+mC^=*tubP-GAL80*^ts^}^10-PEV-80.4^; *Su*(*var*)*3-9*^1^/ *P*{*white*^+mC^=*Act5C-GAL4*}17bF01. A-rfp, A-gfp, B-rfp, and B-gfp are inverted monochrome separations of the RFP and GFP color channels. Images in (A) and (B) are two discs taken from separate individuals and are presented and analyzed to show the common features of the patterns revealed by SwiM System analysis. Pink boxes indicate location of enlarged regions shown in (C-N). **(C-D)** show regions with comparatively low GFP within clones with higher GFP expression, highlighting variation in the timing of GFP activation even within single clones of cells. **(E-H)** show regions with high variation within and between clones; those in (E) and (F) have higher expression than those in (G) and (H). **(I-J)** show single cells without detectable RFP or GFP expression within clones of otherwise-expressing cells, which we interpret as single cells that are not silencing GAL80 expression. **(K-N)** are clones wherein expression of RFP is remarkably variable despite overall similar levels of GFP expression (*cf*. Figure 3C-D).

Although the discs resembled *wild-type* in the number of GFP-expressing cells and clones, the variation of GFP expression was higher than we saw in *wild-type* animals. For example, we observed clones of low GFP expression within encircling clones that had a relatively higher amount of GFP fluorescence (Figure 5C-D). This condition is likely due to a more-recent gain of silencing than in the outlying clones of cells. Moreover, the GFP expression in all GFP-expressing clones of *Su*(*var*)*3-9*^1^ individuals were more heterogeneous in their fluorescence than in *wild-type* (Figure 5E-F). These variations were true whether the expression of GFP in a clone was high on average, or low (Figure 5G-H). The only way we can envision such a situation is, as above in *Su*(*var*)*205*^5^/+ mutants, as extremely-brief periods of silencing producing a dip in GAL80, and activation of RFP and FLP for a short window of time: insufficient to accumulate RFP but sufficient to permanently activate GFP. Some individual cells had undetectable GFP (Figure 5I-J) even while cells in the same clone varied from moderate to bright intensities of GFP fluorescence, leading us to believe that these variations are recent events. We therefore interpret the cell-by-cell variation in these patches as extremely short-lived silencing of GAL80 followed by infrequent genome rearrangement by the few FLP molecules in cells of those clones.

Those clones of cells that did express RFP also exhibited stochasticity and high variation in RFP amount (Figure 5K-L). Cells exhibiting red fluorescence were clonal, retaining the general pattern of PEV, though the clones were generally small and themselves expressed high variation as they did for GFP. In some clones (Figure 5M-N), the individual behavior of cells within clones were especially pronounced, indicating that the cells within the clones no longer were acting in concert. This could arise from a failure to maintain epigenetic memory, or from the level of the Su(var)3-9 HMT being at or near the threshold required for silencing.

Although both *Su*(*var*)*205*^5^ and *Su*(*var*)*3-9*^1^ produce equally derepressed phenotypes as adults, they seem to arrive at that terminal phenotype in different ways. *Su*(*var*)*205* mutants fail to ever establish silencing in most cells. Those few cells in which silencing is established is lost through development, and establishment or reacquisition of silencing is rare or non-existent in mutants. *Su*(*var*)*3-9*^1^ mutants do establish silencing quite well, however they are unable to retain it. Further, once lost, silencing is rarely, if ever, re-established for more than just a brief time. We consider that the *Su*(*var*)*3-9*^1^ mutants may simply express a weaker phenotype than do the *Su*(*var*)*205*^5^ mutants, either because the allele is not amorphic, because other HMTs – such as *G9a* or *eggless* – may partially compensate, or because of *bona fide* separable roles for Su(var)3-9 and HP1a (22).

The presence of any RFP cells in either mutant condition was itself a surprise given the terminal and well-characterized phenotypes in the eyes of adult PEV-expressing flies (*e.g*., on the *white*^mottled-4^ allele (12)) who also bear these mutations. We believe that the ability for any cell to establish silencing, even transiently, must mean that there is at some point sufficient histone methyltransferase activity even though the gene dose has been reduced in *Su*(*var*)*3-9*^1^ mutants. We imagine that the levels of HP1a and HMT fluctuate normally, thus in some cells through development there is ample silencing activity, while in others there is not. In a *wild-type* condition, this leads to PEV phenotypes. In mutants reducing HP1a or HMT, the fluctuations still occur, but at a lower average level below a threshold required to establish heterochromatin-induced gene silencing. Still, a few cells by means of these fluctuations do rise above the threshold and establish silencing but, more often than not and perhaps ultimately inevitably, they lose it again.

### Further Thoughts on Position Effect Variegation and Epigenetic Regulation

Our data from analyzing *wild-type* discs further support our previous conclusion (11), that what we observe as PEV is the final outcome of a life-long series of dynamic switches, from silenced to derepressed and back again. In *Su*(*var*)*205/+* mutants, non-fluorescent cells represent failures to ever establish silencing, accounting for the final fully-suppressed phenotype of these individuals. In contrast, in *Su*(*var*)*3-9/+* mutants, GFP-fluorescent cells represent successful establishment of silencing, but the subsequent near-total loss of that silencing, accounting for its fully-suppressed phenotype.

We suggest that the developmental dynamism in heterochromatin-induced gene silencing we see in *wild-type* or mutant flies modifies the currently-prevailing notion of epigenetic memory in heterochromatin-mediated silencing. In the current understanding, the epigenetic information that accounts for the memory of silencing is the same as the mechanism for silencing – H3K9 methylation and HP1a are thought to be copied to both daughter chromatids in S phase and then enact silencing in the subsequent G1 phase. Our data are not easily reconciled with this view because the losses of silencing we observe (*i.e*., any cell or clone that expresses GFP) would be expected to erase the epigenetic memory.

We envision a few possible reconciliations. It is possible that epigenetic memory and epigenetic silencing are mediated by different and separable mechanisms. For example, histone methylation patterns (by, *e.g*., Su(var)3-9) could be inherited and establish a “proto-silencing” state that itself is not silencing, but when coupled by sufficient nuclear HP1a could then enact silencing (23). In this envisioning, these two factors could be independent with H3K9-methylation accounting for memory and H3K9-methylation+HP1a accounting for the silencing (24). Alternatively, we could instead view heterochromatin-induced gene silencing as requiring constant establishment from neighboring heterochromatin with no epigenetic “maintenance” phase at all through S phase. We prefer this latter view, as it is consistent with others’ work in *Drosophila* and *S. cerevisiae* that have shown that liberation of a variegating gene from juxtaposition to heterochromatin immediately and completely derepresses it (25–26).

Our analysis of mutants that compromise known components of the H3K9 heterochromatin silencing mechanism show that decreased silencing leads to increased fluctuations in instability. We believe that this should be interpreted as a need for constant levels of gene products from both *Su*(*var*)*205* and *Su*(*var*)*3-9* because the functions of those heterochromatin components are themselves dynamic. Dynamism could arise either through transient interactions (16), through competing activities that act to counteract their roles, or from fluctuations in their expression level or activity. One, or all, of these factors may predominate the correlation with silencing. We do not see them as mutually exclusive and, in fact, they may be different aspects of the same mechanism. For example, for the lower concentration of HP1a in mutants, we expect longer times in which H3K9 methylated histones are not bound by HP1a, leading to derepression (16). This may be by simple mass-action (27) or elaborated ideas thereof (10, 28–29), by particular properties of phase transition (30, 31), or otherwise. Similarly, ubiquitously-acting H3K9 demethylases (members of the KDM family) act in opposition to HMT activity. This could explain why an enzyme shows dose-sensitivity, because its activity is in constant competition with other enzymes. While not supported specifically by any experiment of which we are aware, this possibility would account for the results we see with our SwiM analysis.

The SwiM System has allowed us to address some of Dr. Janice Spofford’s questions. We have discovered that PEV is a highly dynamic process, with each gene undergoing repeated rounds of silencing and derepression; we conclude that the degree of reversibility she mused might exist does, and is quite high. More, we do not see evidence of a single time of heterochromatin-induced gene silencing, but instead a constant need to re-establish silencing by neighboring heterochromatin (32). In *Drosophila* and mouse, nucleation of HP1 is capable of recruiting Su(var)3-9, H3K9-methylation, and more HP1a (33–34). However, in these experiments, the silent chromatin created was ephemeral (35), indicating that *bona fide* heterochromatin is able to initiate silencing, and this activity is persistently required. How this acts at genes or promoters near heterochromatin-euchromatin breakpoints is not yet clear, but it is suggestive that the current model that HMT/H3K9/HP1a is sufficient to establish epigenetic memory may be incomplete. For PEV, the manifestation of instabilities or errors in epigenetic memory, it seems evident that there are times in development during which re-establishment is more difficult, and these times correspond to periods more sensitive to genetic mutation and, perhaps, to environmental perturbations.

## Acknowledgements

We gratefully acknowledge support from the National Institutes of Health (R01GM123640). The work was also made possible by support from the National Institutes of Health to the University of Arizona Cancer Center (P30CA023074) and to the Bloomington Drosophila Stock Center (P40OD018537), as well as the University of Indiana Univeristy to the latter. We also acknowledge outstanding resource support from the University of Arizona and the University of Arizona Office of Research, Innovation & Impact for the University of Arizona Core Facilities and the Univeristy of Arizona Cancer Center.

## Materials and Methods

### Strains, husbandry, and genetic crosses

*wild-type* flies were *yellow*^1^ *white*^67c23^. The main SwiM System strain was maintained as *y*^1^ *w*^67c23^; *P*{*white*^+mC^*=tubP-GAL80*^ts^}10-PEV-80.4; *P*{*white*^+mC^=*Act5C-GAL4*}17bF01/TM6B, *Tb*^1^. *P*{*white*^+mC^=*tubP-GAL80*^ts^}10-PEV-80.4 is a transposition of *P*{*white*^+mC^=*tubP-GAL80*^ts^}10 from 51D1 (36) to its current location in the distal centric heterochromatin of chromosome *2* (h35-h36) by exposure to an endogenous source of Δ*2,3 transposase* (*H*{*white*^+mC^=*PDelta2-3*}*HoP8, y w*\*) and selection for strains exhibiting variegation of the *white^+^* gene. For *wild-type* SwiM analysis, male SwiM flies were crossed to virgins of one of two *G-TRACE* strains, either genotype *w*\*; *P*{*white*^+mC^=*UAS-RedStinger*}4, *P*{*white*^+mC^=*UAS-FLP1.D*}JD1, *P*{*white*^+mC^=*Ubi-p63E(FRT.STOP)Stinger*}9F6/CyO or *w*\*; *P*{*white*^+mC^=*UAS-RedStinger*}6, *P*{*white*^+mC^=*UAS-FLP.Exel*}3, *P*{*white*^+mC^=*Ubi-p63E(FRT.STOP)Stinger*}15F2. We saw no difference in these two strains and the choice was based on linkage of the two *Su*(*var*) mutations. *Su*(*var*)*205* was *In*(*1*)*w*^m4^; *Su*(*var*)*205*^5^/*CyO, Cy*, and *Su*(*var*)*3-9* was *In*(*1*)*w*^m4h^; *Su*(*var*)*3-9*^1^/*TM3, Sb*^1^. For mutant SwiM analysis, male SwiM flies were crossed to virgins of genotype *Su*(*var*)*205*^5^/*CyO, Cy; P*{*white*^+mC^=*UAS-RedStinger*}6, *P*{*white*^+mC^=*UAS-FLP.Exel*}3, *P*{*white*^+mC^=*Ubi-p63E(FRT.STOP)Stinger*}15F2 virgins, or *w*\*; *P*{*white*^+mC^=*UAS-RedStinger*}4, *P*{*white*^+mC^=*UAS-FLP1.D*}JD1, *P*{*white*^+mC^=*Ubi-p63E(FRT.STOP)Stinger*}9F6/*CyO*; *Su*(*var*)*3-9*^1^/*TM3, Sb*^1^.

For all of the above strains, we created isogenic strains at the outset of this work by crossing individual males to a double-balancer strain (*y*^1^ *w*^67c23^; *wg*^Sp-1^/*CyO*, *Cy*; *Pr*^1^ *Bsb*^1^ *ry*^506^/ *TM6B, Tb*^1^ *Hu*^1^). Individual Curly Tubby male offspring were backcrossed to individual double-balancer females, then bred *inter se* to produce strains derived from single chromosomes (*X* by means of single paternity, chromosomes *2* and *3* by single paternity and balancers and mitochondrial genomes by single maternity; chromosome *4* was not intentionally isogenized). These individually isogenic strains differ from each other, but individual flies within each strain has no or limited polymorphism. Siblings for the experiments shown in Figures 2, 3, 4, and 5 are genetically identical to each other as they are from genetic crosses between isogenized strains.

Flies were maintained in glass vials, fed “Karpen-Eckert” medium (30 g/L yeast extract, 55 g/L corn meal, 11.5 g/L agar, 72 mL/L dark molasses, 6 mL/L propionic acid, 2.4 g/mL tegosept), and raised at 25°C at 80% humidity; manipulation was done after etherization.

### Photography

Images of whole flies were taken with a Sony a7iii camera back attached to a Nikon SMZ-1500 microscope, illuminated with a Peak Plus Tactical LED Flashlight. Flies were cradled in a 90° angle constructed of two mirrors to simultaneously image both eyes. Pigment categories in Figure 2 were determined by photographing the eyes, quantifying percent coverage using ImageJ, and binning the categories (here into pentiles), as we have done before (11, 37–39).

### Dissection, Microscopy, and Fluorescence Detection

Larval imaginal discs were dissected from wandering third instar larvae in 1X PBS, and were visualized and photographed using a Zeiss AxioZoom.v16 equipped with series 00 (excitation BP 530-585, beamsplitter FT 600, emission LT 615), 38HE (excitation BP 470/40, beamsplitter FT 495, emission BP 525/50), or 74HE (excitation DBP 480/30 + 565/25, beamsplitter DFT 505 + 583, emission DBP 525/31 + 616/57) filter sets. Fluorescence quantification in Figure 2E was done by integrating fluorescence intensity over the entire disc using the Zeiss ZEN (version 2.3-blue) software. Those data were analyzed for goodness-of-fit using a non-parametric chi-square test because they are categorical data. Regression was done using Kendall’s robust line-fit method according to (40). Images presented here were processed for bright-contrast using the “Best” algorithm of the same software before export as JPEG. Black-on-white separations were made in Adobe Photoshop CC (version 20.0.4) and images were cropped using Photoshop or Apple Pages (version 11.2). Bright-contrast was only altered where indicated (by an asterisk in the figure), decreasing brightness and increasing contrast maximally or near-maximally to highlight and make visible very low-levels of expression; expression levels (absolute or relative) are not interpretable in images manipulated in this way.

